# Resolving a QTL complex for height, heading, and grain yield on chromosome 3A in bread wheat

**DOI:** 10.1101/2020.02.14.947846

**Authors:** Alba Farre Martinez, Clare Lister, Sue Freeman, Jun Ma, Simon Berry, Luzie Wingen, Simon Griffiths

**Author notes:** These authors contributed equally to this work (Alba Farre Martinez, Clare Lister).

## Abstract

Crop height (Ht), heading date (Hd), and grain yield (GY) are interrelated traits in wheat. Independent manipulation of each is important for adaptation and performance. Validated QTL for all three collocate on chromosome 3A in the Avalon x Cadenza population. We asked if these are linked or pleiotropic effects. The region was dissected using recombinants derived from Near Isogenic Lines. It was shown that Ht and Hd are controlled by independent genes. The newly defined Ht QTL interval contained a gene cluster involved in cell wall growth and displaying high levels of differential transcript expression. The Hd locus is much larger and rearranged compared to the reference genome but *FT2* is a candidate of particular interest. The Hd effect was shown to act independently of photoperiod and vernalization but did exhibit genotype x environment interaction suggesting a role in ambient temperature sensitivity. It was the Hd locus that was most associated with increased GY of Cadenza alleles, supporting physiological studies proposing that ‘late’ alleles at this locus increase spike fertility and grain number. The work has uncoupled height from heading and yield and shown that one of very few validated GY QTL in wheat is probably mediated by phenological variation.

**Highlight:** There only are three validated wheat yield QTL. Here, one of them was genetically dissected.

This showed that the physiological basis of the yield effect is likely to be phenological.

## Introduction

Breeding for increased grain yield (GY) is achieved in concert with the optimisation of other important agronomic traits, notably height (Ht) and heading date (Hd). While the direction of selection for GY is always for increase, the other two traits are adaptive which means that the optimum Ht and Hd of the crop depends very much on the target environment. This presents a challenge because all three traits are often related. For example, selection for yield alone can result in tall (Law *et al.*, 1978) late heading lines (Bogard *et al.*, 2011) which are prone to lodging, late season stress, and delayed harvest.

A small number of major genes have been used by breeders to mould their germplasm into primary adaptive classes targeting broad environmental categories such as those captured in the mega-environment concept (Rajaram *et al.*, 1993). So, for Ht major semi-dwarfing genes are deployed such as *Rht-B1b, Rht-D1b* (Miralles and Slafer, 1995) and *Rht8* (Worland *et al.*, 1998b). For Hd, photoperiod and vernalisation sensitivity genes are used in a similar way with the consequence that allelic classes for groups of major genes are often fixed, or at least represented at a high frequency, within germplasm pools targeting specific mega-environments (Kiss *et al.*, 2014). For photoperiod response these are mainly homoeoalleles of *Ppd-1* (Worland *et al.*, 1998a), *Ppd-2* (Zikhali *et al.*, 2017), and for vernalization, mainly *Vrn-1* (Iwaki *et al.*, 2001),

After these major gene combinations have been optimised, genetic progress for adaptation, and its interplay with grain yield, made within these pools depends on polygenic variation comprising individual effects which are relatively small and identified as quantitative trait loci (QTL). More detailed investigation into these effects, often relying on their conversion to discrete Mendelian factors by the production of Near Isogenic Lines (NILs), facilitates a deeper understanding of the function/s of gene/s underlying the QTL which are being used by breeders in that programme. In this NILs development facilitated the characterization of a heading date QTL identified on 1DL as an earliness *per se* (*eps*) effect (Zikhali *et al.*, 2014) which was then described as a discrete Mendelian factor called *EPS-D1* caused by deletion of *ELF3* (Zikhali et al., 2015). Analysis of floral development showed that the late alleles of *EPS-D1* increased spike fertility through enhanced floret survival (Prieto *et al.*, 2018). Controlled environments were used to show that *EPS-D1* displayed a strong interaction with ambient temperature variation (Ochagavía *et al.*, 2019). A similar depth of understanding of all major QTL segregating in elite germplasm pools is an important foundation stone for future genetic gains in genomics led breeding programmes.

Through the use of segregating populations or association panels derived from varieties which are well adapted to the same environment a number of studies have been able to identify the QTL that are being used to achieve incremental yield gains and fine tuning of adaptation. The Avalon x Cadenza doubled haploid (DH) population was used by us as part of a meta QTL study to describe variation in UK wheat for Hd (Griffiths *et al.*, 2009), Ht (Griffiths *et al.*, 2012) and GY (Ma *et al.*, 2015). A library of NILs was then produced in which reciprocal transfers of Avalon and Cadenza alleles into the opposing variety allowed QTL validation in both parental contexts (Farré *et al.*, 2016). This allowed us to validate the original Avalon x Cadenza QTL and prioritise key effects. A QTL identified on chromosome 3A was of particular interest as the same locus affected GY, Ht, and Hd with the Cadenza allele increasing all three. Very few robust GY QTL have been identified in wheat and an even smaller number validated by comparison of NILs. These include: a grain size effect on chromosome 6A that increases GY (Simmonds *et al.*, 2014) and *Rht-1* semi dwarfing increases yield via increased grain number in some environments (Miralles and Slafer, 1995). The Avalon x Cadenza 3A GY QTL and derived NILs increase GY through grain number (Farré *et al.*, 2016). This is important because almost all of the genetic gains achieved for GY and GY plasticity in wheat have been as a consequence of increased grain number (Slafer *et al.*, 2014).

It was not known whether the Hd and Ht effects collocated with GY at the 3A locus are associated by genetic linkage or pleiotropy. The aim of this study is to show whether they are genetically distinct, increase understanding of the mechanism to show how they might contribute to adaptation and GY, and develop assays facilitating precise marker assisted selection at this locus.

## Materials and methods

### Development of NILs and recombinants

The A × C DH population was one of several developed to represent a broad spectrum of the variation present in the UK elite winter germplasm pool and is now the UK reference population under the UK Department of Environment, Food and Rural Affairs (DEFRA) Wheat Genetic Improvement Network (WGIN). Several Hd, Ht, and GY QTLs have been previously identified in the A × C DH population (Griffiths *et al.*, 2009; Griffiths *et al.*, 2012; Ma *et al.*, 2015). In this work we are focused on the Hd, Ht, and GY QTL located on chromosome 3AS. Both Avalon (UK winter wheat) and Cadenza carry photoperiod sensitive alleles of *Ppd-D1* and *Ppd-B1*. Avalon carries the winter alleles of *vrn-A1, vrn-B1* and *vrn-D1* whereas Cadenza carries a dominant *Vrn-A1a* allele conferring a facultative/spring growth habit.

The development of families of Near Isogenic Lines (NILs) and their use for the validation of the 3A Hd, Ht, and GY QTLs is described in (Farré *et al.*, 2016). For the development of recombinants within the introgressed segment BC_2_ (BC_3_ equivalent because the backcross donor parent was a line from the Avalon x Cadenza segregating doubled haploid population) heterozygous plants were self-pollinated. Lines heterozygous for a region between simple sequence repeat (SSR) markers *wmc505* and *wmc264*, identified as a meta-QTL region for flowering on 3A in Griffiths et al (2009), were self-pollinated to generate a BC_2_F_3_ of 454 individuals. These markers were then used to identify recombinants in this interval. Eighty four of these BC_2_F_3_ recombinants were self-pollinated and homozygous BC_2_F_4_ recombinants selected using the two flanking SSR markers. A total of 76 recombinant BC_2_F_4_ lines were selected for further analysis.

### Assessing photoperiod sensitivity of 3A heading date effect

The A × C BC_2_ NILs were grown under controlled environments. Seeds were sown in January 2014 and grown in an unheated but daylength-controlled glasshouse and therefore fully vernalized at 6-10°C using natural vernalization, under short days (SD, 10h light) for eight weeks. The plants were then grown at 13-18°C under two photoperiod treatments SD or long days (LD, 16h light)). Plants under SD and LD were grown with natural light for 10h and the LD plants had an additional artificially extended photoperiod of 6h using tungsten bulbs. We used eight 60W tungsten lamps spaced 90 cm apart and 2.1m above the plants; delivering 1 micromole s^-1^ m^-2^. The plants were grown in a randomized complete block design with three replicates. NILs were classified according to their genotype across the *wmc505* – *wmc264* genetic interval (twenty-four with the Avalon and nineteen with the Cadenza introgression). To verify that the NILs had been adequately vernalized, five plants each of the winter wheat cultivars Claire, Malacca and Hereward were grown as controls. Hereward flowers more than 30 days later than Malacca and Claire when incompletely vernalized for four weeks and this was associated with copy number variation at *Vrn-A1* (Diaz et al., 2012). Days to ear emergence (Hd) was scored as the number of days after the 28th of April when the ear was more than 50% emerged from the flag leaf on the main shoot and corresponding to Zadoks stage 55 (Zadoks *et al.*, 1974). Data was evaluated using two-way ANOVA in which the interaction between treatment and allele was included in the model. ANOVA was performed using Genstat 16^th^ edition (VSN International).

### Assessing vernalization sensitivity of 3A heading date effect

For the controlled environment experiments, 32 out of 76 chromosome 3A recombinants were selected based on the extent of the Avalon introgression. The plants were grown under different vernalization treatments (0, 4, 6 and 8 weeks under SD at 6°C) and then transfered to LD photoperiod (as described above). The plants were grown in a randomized complete block design with three replicates for each treatment. The mixed model used included treatment, allele and the interaction between treatment and allele as fixed factors, and blocks as a random factor. Blocks were considered random factors in the model in order to recover inter-block information due to the presence of missing values. The mixed model was fitted using linear mixed procedures from Genstat 16^th^ edition (VSN International). Heading date was recorded as days to ear emergence (Hd). In this case, Hd indicates the difference between days from 50% ear emergence and the day that plants were transferred from the vernalization treatment to LD. The winter wheat cultivars Claire, Malacca and Hereward were also grown under these conditions as controls.

### Measurement of developmental phases

The NILs (NIL-A and NIL-C) were used to determine which developmental phases were affected by the 3A QTL. NIL-A (carrying the Avalon introgression in the QTL region) and NIL-C (carrying the Cadenza introgression) came from the AC179-E27-2 stream (Farré *et al.*, 2016). Plants were vernalized for 8 weeks at 6°C under SD and then transferred to a glasshouse with a temperature around 18°C and LD photoperiod. Apices were dissected from three randomly selected plants of each NIL every 2-3 days and examined under a light microscope. This allowed the time from sowing to double ridge (S-DR), double ridge to terminal spikelet (DR-TS) and from terminal spikelet to heading (TS-HD) to be determined, following the scales proposed by (Kirby and Appleyard, 1987). Hd was scored as above.

### Statistical analysis of GY

A simple linear model (function lm in R vs. 3.6.1) was fitted to analyse the relationship between a trait and the genetic markers in the QTL region. P-values were calculated from the t-statistics. Box-plots were plotted using function boxplot in R.

## High resolution mapping of the 3A Hd and Ht QTLs

### Phenotype evaluation

Seventy six recombinant BC_2_F_4_ lines were phenotyped in two field experiments (spring-sown and autumn-sown) and under controlled environments. Field trials were conducted at Church Farm, Bawburgh, Norfolk, UK, in 2013 (spring and autumn sown). Details of meteorological conditions are given in Supplementary Fig. S1. Experimental design followed a randomized complete block design with three replicates. Plots consisted of 4 rows, 1m long and 12cm apart and grown according to standard agricultural practice, except that plant growth regulators (PGRs) were not applied. The trial included both parents (Avalon and Cadenza) and NIL-A and NIL-C in each replicate. Hd was assessed in thermal time (°C d, using a base temperature of 0°C). Ht was measured from soil level to the tip of the ear (cm). None of the material was awned so this is not a complicating feature in the description of height.

### Genetic mapping and QTL analysis

Genomic DNA extraction was performed using published protocols (http://maswheat.ucdavis.edu/PDF/DNA0003.pdf), adapted from Pallota et al. (2003). To increase marker resolution across the 3A QTL region, 65 additional markers were chosen. These were mainly KASP markers selected from the integrated 3A genetic map at CerealsDB (http://www.cerealsdb.uk.net/cerealgenomics/CerealsDB/kasp_mapped_snps.php) or KASP markers derived from iSelect markers (Avni *et al.*, 2014). Marker information can be found at CerealsDB (http://www.cerealsdb.uk.net/cerealgenomics/CerealsDB/iselect_mapped_snps.php).

Methods for genotyping with SSR and KASP assays used have been described previously in (Wingen *et al.*, 2014) and (Zikhali *et al.*, 2014) with the precise conditions used dependant on the specific primer pairs. Linkage analysis was performed using JoinMap® version 3.0 (Ooijen and Voorrips, 2002), using the default settings. Linkage groups were determined using LOD threshold of 3.0 and genetic distances were computed using the Haldane mapping function. Genstat 16^th^ edition was used for QTL detection and to estimate QTL effects using single marker analysis and the composite interval mapping (CIM) function.

### Identification and sequencing of candidate genes

Prior to the publication of the IWGSC RefSeq v1.0 genome assembly (Appels *et al.*, 2018) a variety of sources were used to identify candidate genes related to flowering or development within the QTL regions, from the available rice, *Brachypodium*, barley and wheat sequences. Primers were designed to amplify the A-genome homoeologue of these candidates. PCR and sequencing reactions were carried out following the methods described by (Zikhali *et al.*, 2014). Newly discovered SNPs between Avalon and Cadenza were converted to KASP markers using PolyMarker (Ramirez-Gonzalez *et al.*, 2015) and validated in the BC_2_F_4_ recombinant lines. With the publication of IWGSC RefSeq v1.0 genome assembly it was possible to scrutinise all the genes covering the Ht and Hd QTLs from the sequence of Chinese Spring, with a region of 3AS between 45-210 Mb analysed. In addition the RNAseq data (see below) was visualized in the Integrative Genomics Viewer (IGV, https://software.broadinstitute.org/software/igv) to identify SNPs. This approach identified an additional cohort of candidate genes from which new KASP markers were generated as above, where possible.

### RNAseq BSA strategy

Three recombinants from either the extreme early individuals (Avalon allele) or late flowering (Cadenza allele) individuals were combined into two bulks. The bulks also carried the Avalon or Cadenza alleles, respectively, for height. Plants were vernalized for 8 weeks at 6°C under SD and then transferred to the glasshouse with a temperature of around 18°C and LD photoperiod. The time-point for sample collection, between double ridge and terminal spikelet, was selected based on the results of the developmental phase experiment. Total RNA was prepared from the whole plant of each recombinant separately using the RNeasy^®^ Plant Mini Kit (Qiagen), followed by treatment with DNase I using the RNase-Free DNase Set (Qiagen). RNA purification was performed using the RNeasy^®^ kit (Qiagen), according to the manufacturer’s protocol. Equivalent amounts of RNA from the three early or late recombinants were mixed to produce each RNA bulk sample.

Library construction and sequencing was performed by The Genome Analysis Centre (TGAC) in Norwich, UK. One Illumina TruSeq RNA version 2 library was constructed per bulk. Sequencing was carried out on the Illumina HiSeq2000 with 100bp paired-end reads. The resulting reads were mapped to the wheat reference sequence using the RNAseq aligner STAR, with all default parameters chosen. The resulting BAM files were then processed with SAM tools and analysed using default parameters in Cufflinks, with reads being mapped to the reference sequence in FASTA format and reference annotation in GFF3 format. The Cufflinks output contained FPKM values for each bulk sample. RPKM values were normalised by setting an RPKM cutoff in order to eliminate false discovery of high fold-changes between genes with very low absolute expression levels (<0.1). Additionally, genes with an expression of 0 were rounded to a small number (0.001) to avoid logarithms of zero. Expression values were then log2 transformed and genes were defined as differentially expressed if they showed an absolute fold-change > 2.

### Yield effects in selected 3A recombinants

Eight recombinant lines, together with parental (Avalon and Cadenza) and NIL (NIL-A and NIL-C) controls, were chosen to assess GY effects. They were sown in 6 m^2^ plots at a seed density of 250 per m^2^ in October 2015 (Morley Farm) and October 2016 (Church Farm) in three replicates, in a fully randomised block design. GY was measured using a combine harvester and thousand grain weight measured using a Marvin seed analyser (GTA Sensorik). Crop height and heading date were scored as described above.

## Results

### Environmental sensitivity of 3A Hd and Ht QTL

The effect of the 3A Hd (Griffiths *et al.*, 2009), Ht (Griffiths *et al.*, 2012), and GY (Ma *et al.*, 2015) QTL has been validated using the same isogenic materials under UK field conditions (Farré *et al.*, 2016). It is important to understand whether the 3A phenology effects are conditioned by sensitivity to photoperiod or vernalization. BC_2_ NILs segregating for the QTL region between *wmc505* and *wmc264* were assessed for Hd under controlled environments with fixed photoperiods and after a saturating vernalization treatment. In the photoperiod experiments, the Avalon 3A NILs headed significantly earlier then the Cadenza NILs (*P*<0.001) (1.58 and 2.48 days under long day (LD) and short day (SD) conditions, respectively, Fig. 1A) without any significant interaction (*P*=0.321; interaction treatment*allele) which shows that the 3A heading QTL is not a photoperiod sensitivity (*Ppd*) effect.

**Fig. 1.**
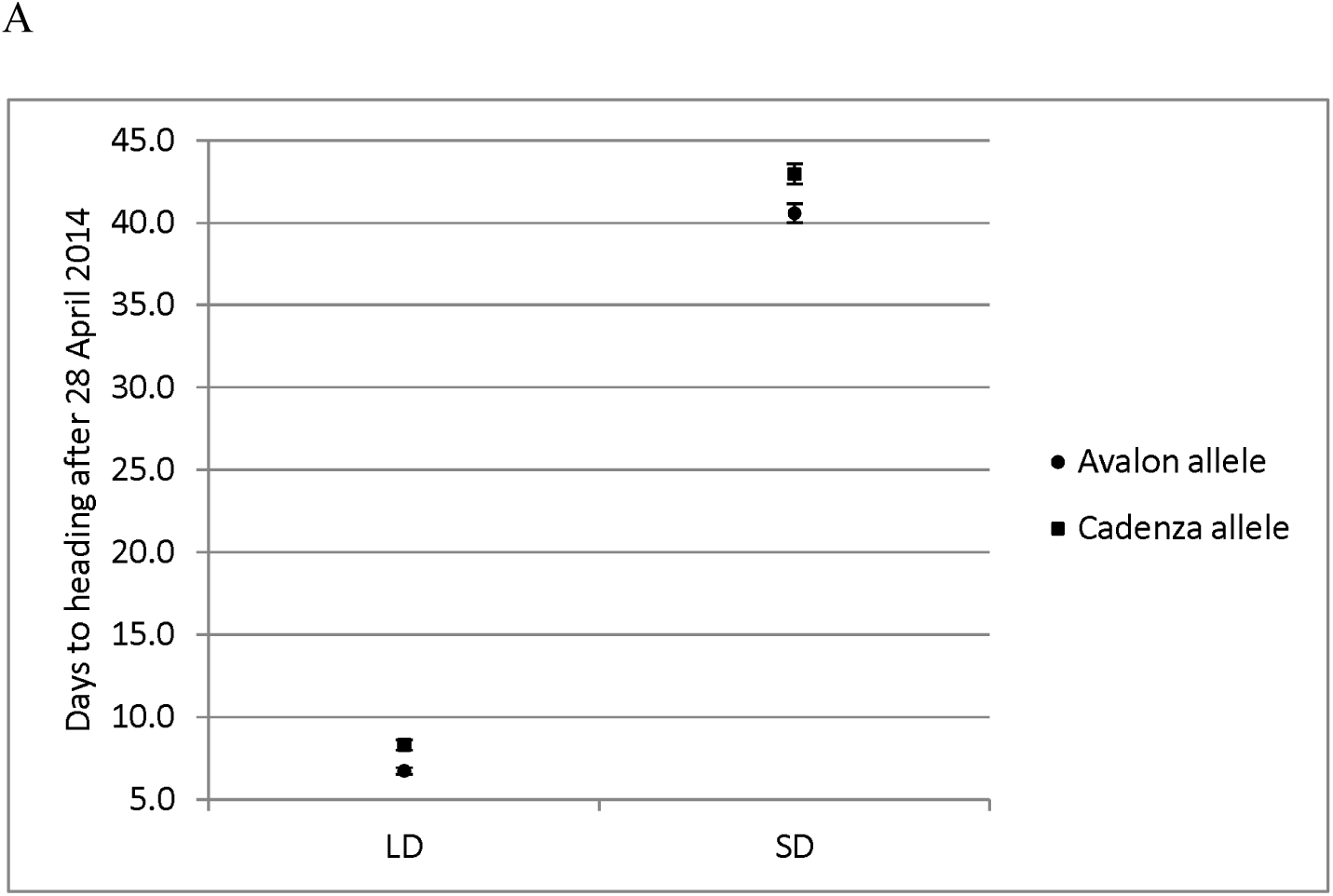
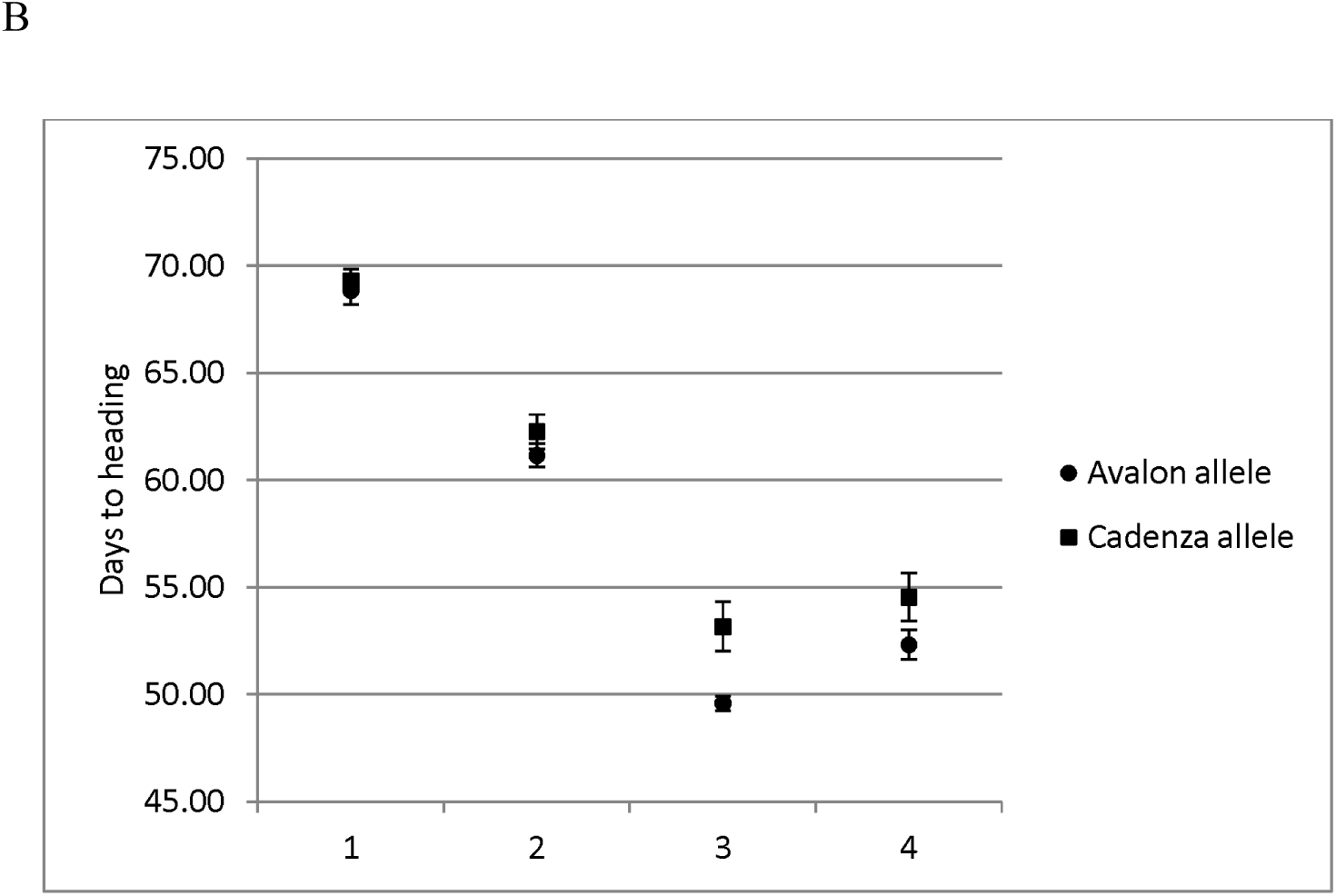
Average days to ear emergence (G55) for leading tillers of controlled environment grown Avalon x Cadenza NILs. ● and ▪ represent the NILs carrying the Avalon and Cadenza alleles at 3A, respectively. A) Short days (SD) and long days (LD) corresponds to 10 and 16 hrs light, respectively. B) response to vernalization using selected recombinant lines. Treatments of 0, 4, 6, and 8 weeks. The vertical bars indicate standard deviation.

To quantify vernalization response 32 out of the 76 recombinants used for fine mapping were selected to be grown in controlled environments (under LD) after vernalization treatments of 0, 4, 6 or 8 weeks. The lines carrying the Avalon allele flowered earlier than those with Cadenza allele for all vernalization treatments (Fig. 1B). Overall a saturating vernalization treatment reduced heading date by around 2 weeks; Cadenza has facultative growth habit and carries the dominant *VrnA1a* spring allele. Based on Wald testing the allelic effect on heading date was highly significant (p=<0.001) but there was no significant interaction between allele and vernalization treatment (p=0.192). However, the mean difference in heading date does increase with a longer duration of vernalization up six weeks. From this we conclude the 3A Hd QTL is not involved in vernalization sensitivity and, taken together with our day length response data, confirms the designation of the 3A Hd effect as earliness *per se* (*eps*) which is most strongly expressed in fully vernalized plants.

### 3A eps QTL affects the duration of early developmental phases

To determine which developmental phases were affected by the 3A *eps* gene, the time from sowing to heading was divided into three phases: from sowing (S) to double ridge (DR), from DR to terminal spikelet (TS), and from TS to heading (Hd). NIL-A (carrying the Avalon allele) and NIL-C (carrying the Cadenza allele) are a pair of NILs from the AC179-E27-2 stream. The differences between them were two and one days from S to DR and DR to TS, respectively. No differences were detected between the NILs in the stem elongation period from TS to Hd (Figure 2, 3). These results indicate that the *eps* region affects the vegetative and early reproductive phases.

**Fig. 2.**
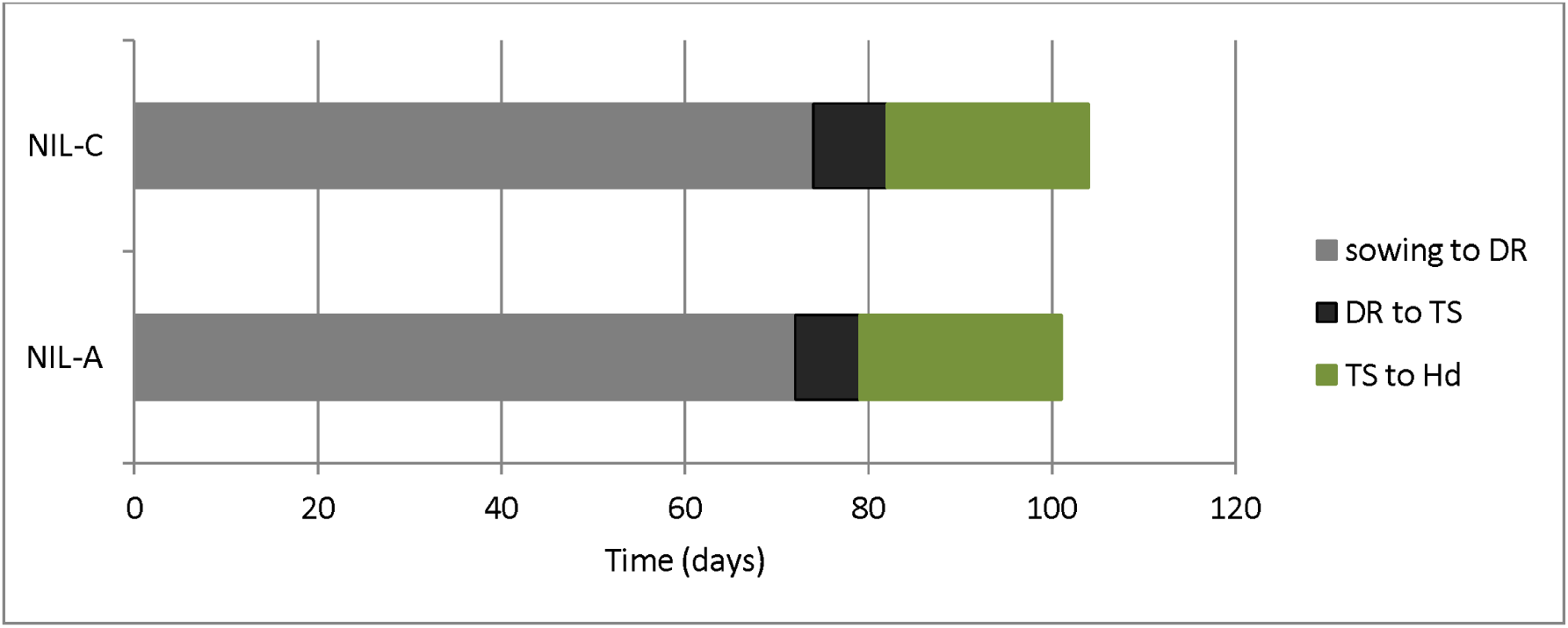
Duration of developmental phases of NIL-A (carrying the Avalon allele) and NIL-C (carrying the Cadenza allele). Developmental phases were divided into three phases: from sowing to double ridge (DR) in grey, from DR to terminal spikelet (TS) in black, and from TS to heading (Hd) in green.

**Fig. 3.**
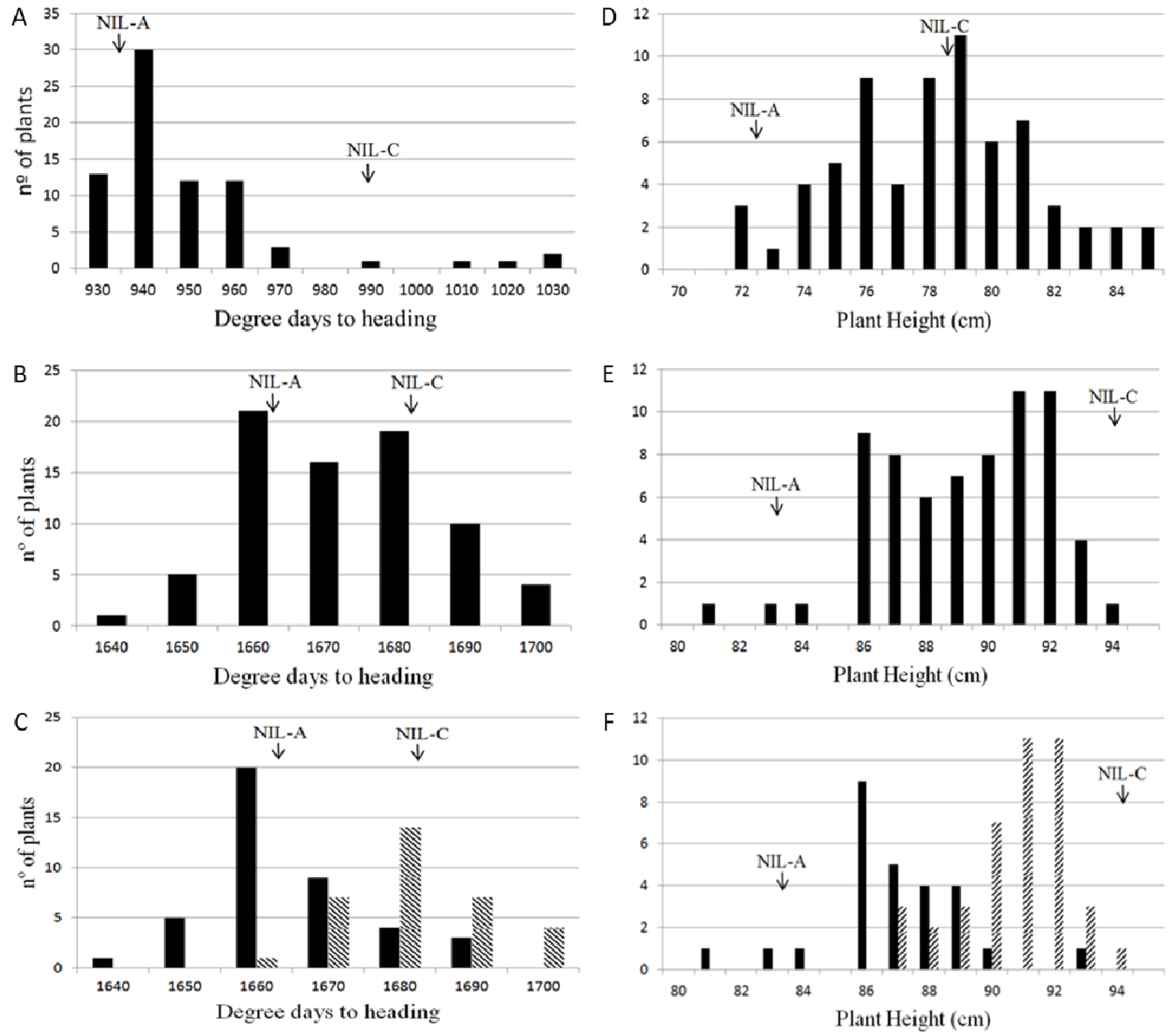
Phenotypic frequency distribution of degree-days-to-heading and plant height in the BC_2_F_4_ population consisting of 76 lines in (A, D) spring-sown and (B, E) autumn-sown. Arrows indicate means of degree days to heading and plant height for NIL-A and NIL-C. (C, F) bars indicate lines with two genotype classes: Homozygous for Avalon (black) and Cadenza (shaded) alleles according to the allele at the wmc505 and BS0003801 locus for Hd and Ht, respectively, using the autumn-sown data.

## High resolution mapping of the 3A Hd and Ht QTLs

### Phenotype evaluation of recombinants

In order to compare the segregation of Ht and Hd at the 3A locus, 76 recombinant BC_2_F_4_ lines derived from crosses of NIL-A and NIL-C with Paragon were phenotyped in two field experiments (one spring-sown and one autumn-sown; Figure 3) and under controlled environments (Figure 1). For both sowing dates in natural field conditions, NIL-A (carrying Avalon allele) flowered earlier than NIL-C (with Cadenza alleles) (938.78 and 1663 mean degree days to heading for NIL-A and 990.0 and 1681 mean degree days to heading for NIL-C in spring and autumn-sown, respectively). However, only the autumn-sown experiment showed a significant difference (p-value=0.017) in mean degree days to heading of the NILs for the 3AS QTL. In contrast, the height difference between NILs was significant in all experiments.

### Mapping and QTL analysis

In order to refine the genetic interval containing the Hd, Ht, and ultimately GY effects, the NIL parents and 76 recombinant BC_2_F_4_ lines were screened using 65 KASP markers. These data were used to perform QTL analysis for the BC_2_F_4_ population for heading date and height to detect the QTLs. The results are presented in Figure 4. Significant Hd and Ht QTL were detected in the 3A region. The peak marker for Hd was BS00021976, which showed a positive effect for the Cadenza allele compared to the Avalon allele (additive effect=1.2 days); this was only observed in the autumn-sown trial. As expected from the NIL data, earlier flowering time is associated with the Avalon allele. The 3A Hd QTL contributed 36.15% of the phenotypic variation in heading date under field conditions. Ht is controlled by an independent QTL with the peak marker BS00022844 (which co-segregates with BS00003801, Fig 5), again the allelic direction is the same as for the NILs with Cadenza increasing.

**Fig. 4.**
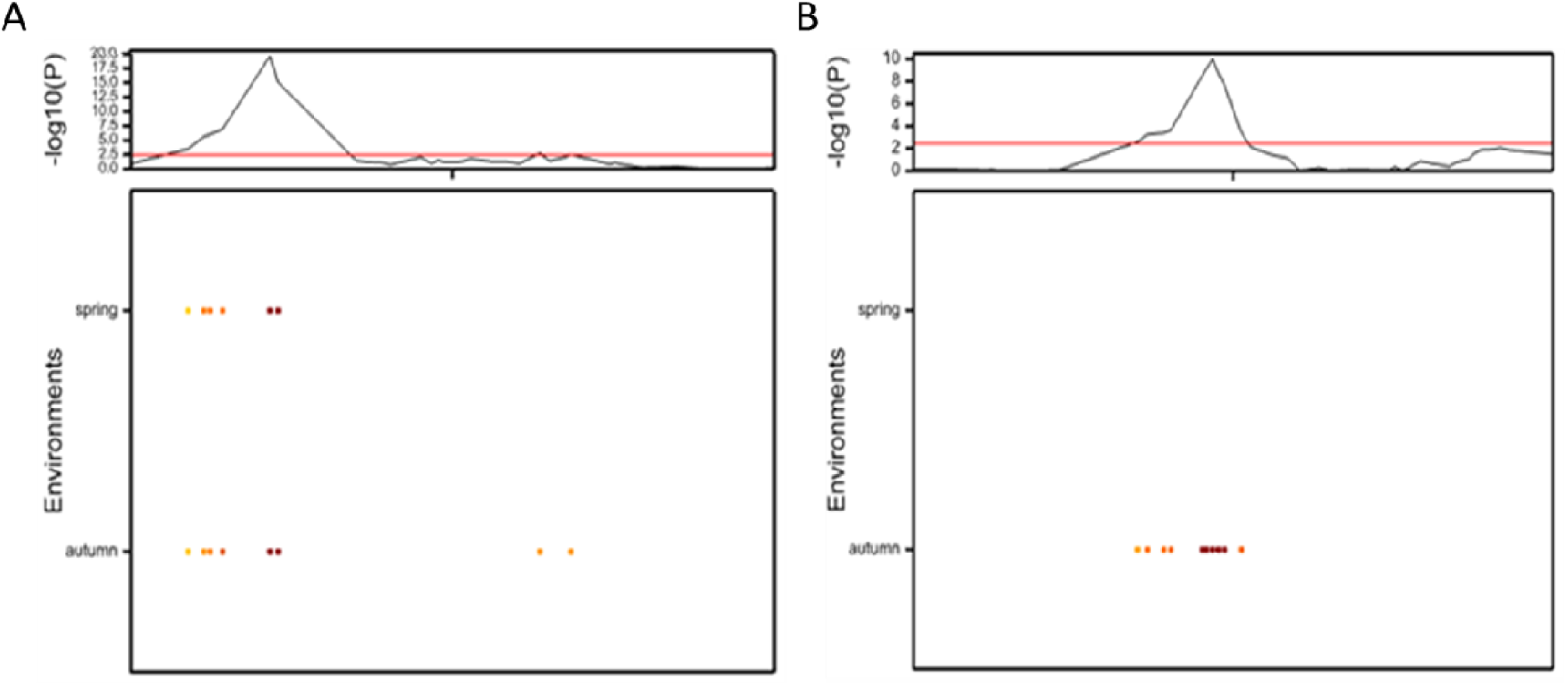
Presence of a QTL for Ht (a) and Hd (b) on chromosome 3A using the data from the 76 BC_2_F_4_ population for two trials, spring and autumn sown. Top panel shows the genome-wide profile, the black and red lines indicate the profile of –log10 (P-value) for a composite interval mapping scan. The red horizontal line shows the threshold value for significance (LOD=2.53). Below each graph is a representation of QTL additive effects detected from spring and autumn sown experiments. The darker orange/brown colours indicate increasing additive effect with the Cadenza allele increasing for Ht and Hd.

**Fig 5.**
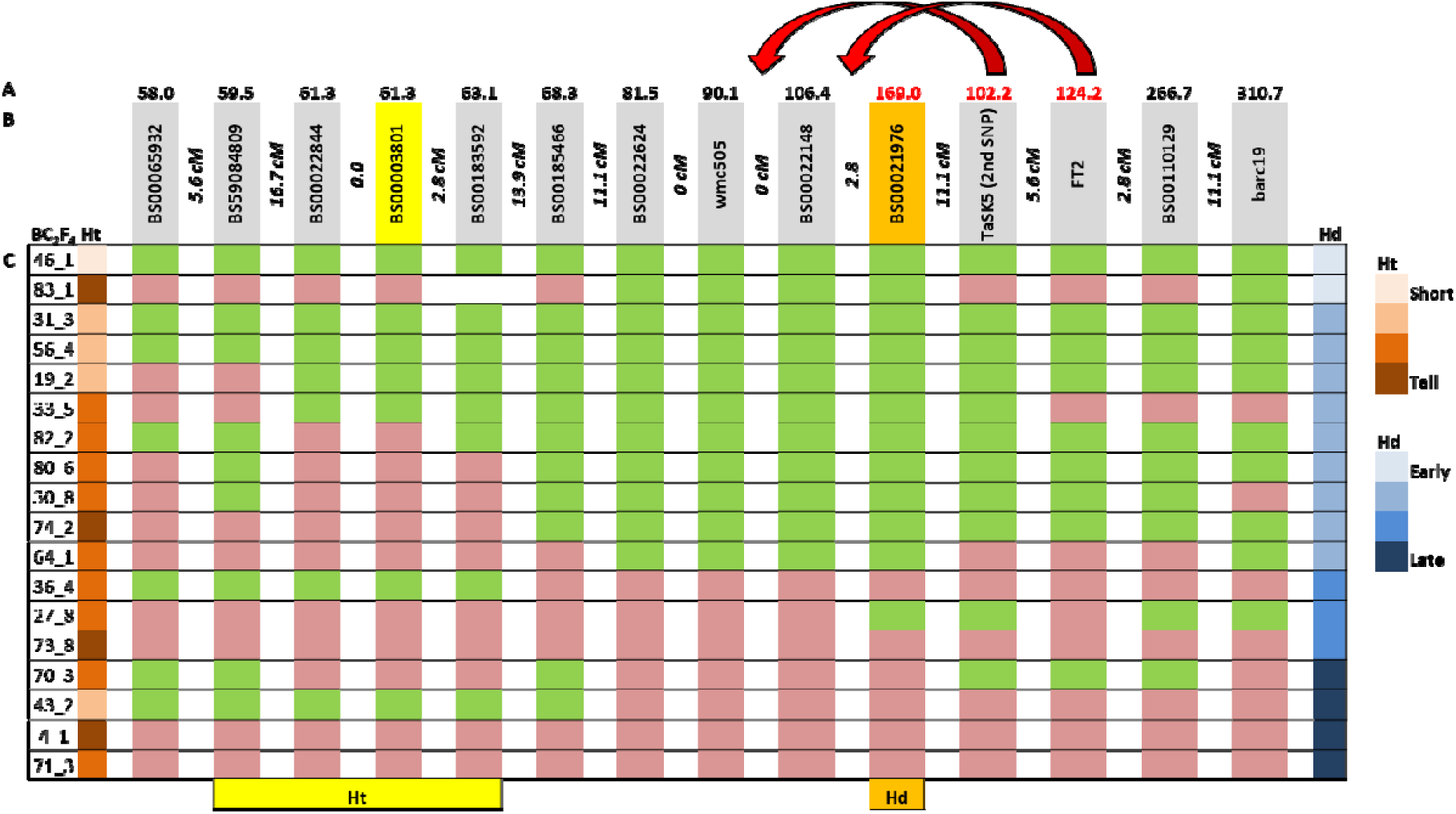
High resolution genetic map and physical map of the Ht and Hd QTL regions on chromosome 3A. A. Position of markers in IWGSC RefSeq v1.0 genome assembly from Chinese Spring. Arrows indicate the suggested rearrangement. B. Markers and genetic intervals between markers. The peak marker for Ht indicated in yellow and the peak marker for Hd indicated in orange. C. Genotyping data from 18 NILs. Green = Avalon at marker, pink = Cadenza, and white = missing data. Location of the Ht and Hd QTLs indicated by yellow and orange blocks, respectively.

Sorting a subset of the recombinants according to Hd (early to late) from the autumn-sown trial, the region containing the 3A *eps* QTL was defined of 13 cM around BS00021976 (Figure 5).

### Examination of gene candidates

Publication of the IWGSC RefSeq v1.0 genome assembly from Chinese Spring (Appels et al., 2018) allowed accurate positioning of the markers defining the QTL loci on Chr 3AS. The closest marker to the Ht QTL is BS00003801 at approximately 60 Mb in the IWGSC RefSeq v1.0, and to the Hd QTL is BS00021976 (at approximately 169 Mb). The genomic sequences from the regions flanking these markers, approximately 5 Mb flanking BS00003801 and 60 Mb flanking BS00021976, were analysed for gene content. The sequences of the unspliced transcripts in these two regions were obtained using Biomart (EnsemblPlants) and then used to search the non-redundant protein sequences database for higher plants with blastx using the default parameters, at NCBI. This produced a list of 146 genes, for Ht, and 536 genes for Hd, with approximately 75% assigned a gene identity. There are at least 10 plausible gene candidates for each QTL, shown in Tables 1 and 2., based on gene function and/or expression level from the RNAseq analysis. Supplementary Table S1 shows the full gene content for the Ht and Hd regions.

**Table 1.**
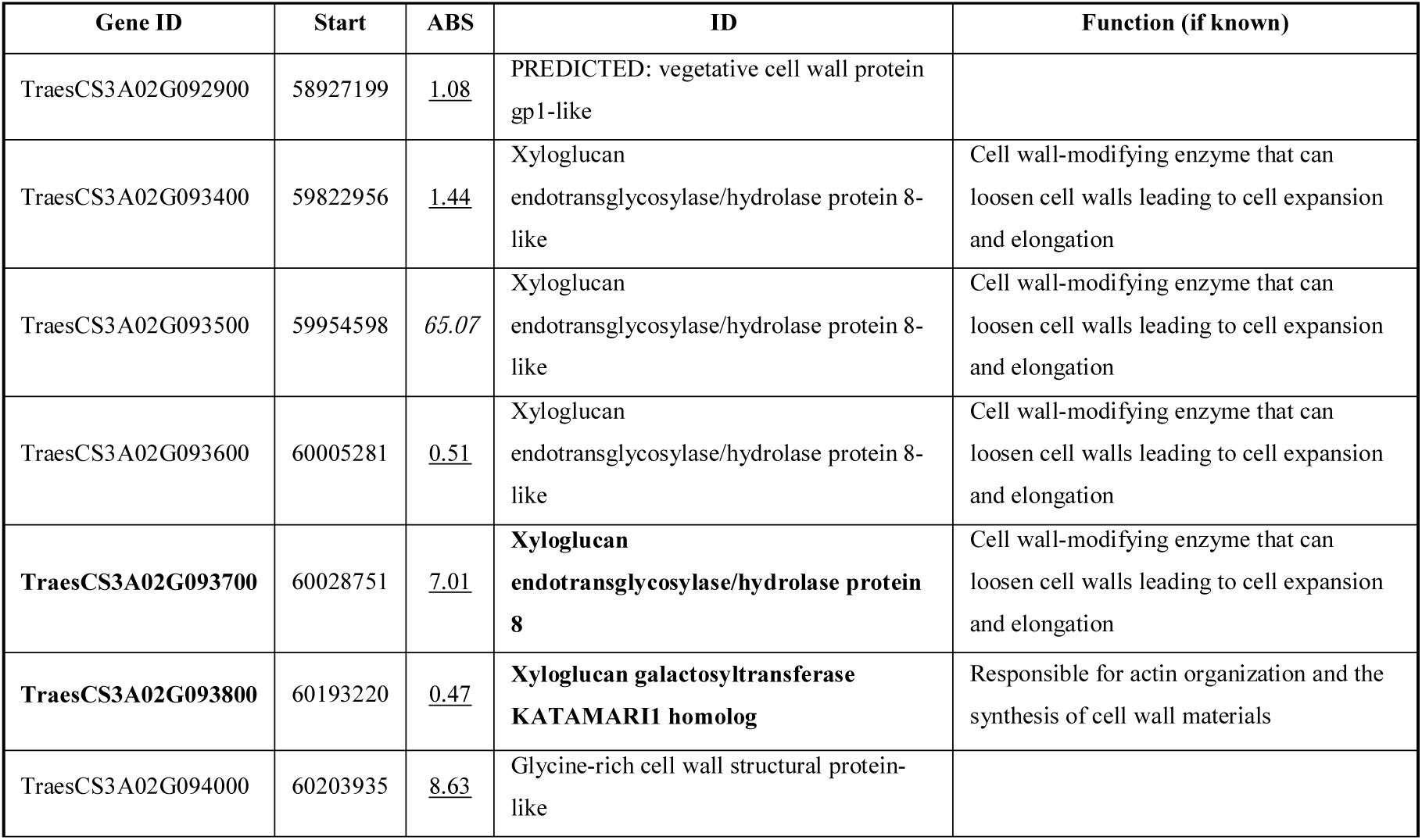
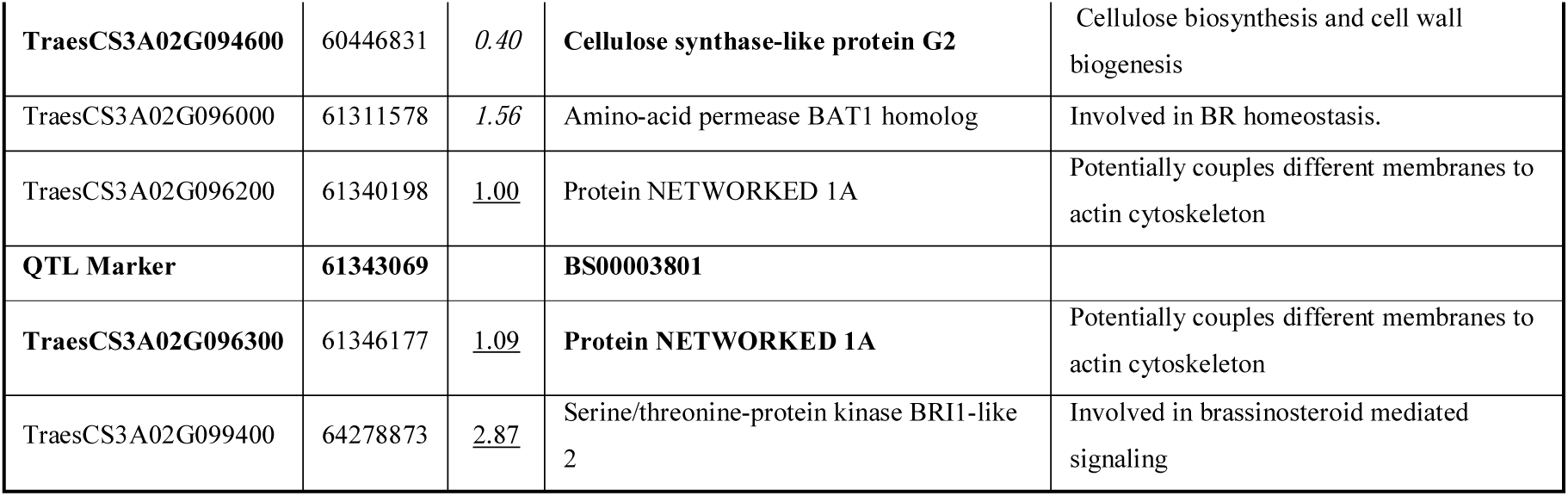
List of candidate genes around the marker closest to the Ht QTL, BS00003801. Transcript name, start position (in Chinese Spring refrence sequence), expression level ratio between Avalon and Cadenza (ABS, <u>higher value</u> from Avalon, *higher value* from Cadenza), gene identity and gene function, if known. Genes for which KASP markers have been developed are shown in bold.

| Gene ID | Start | ABS | ID | Function (if known) |
| --- | --- | --- | --- | --- |
| TraesCS3A02G092900 | 58927199 | <u>1.08</u> | PREDICTED: vegetative cell wall protein gp1-like |  |
| TraesCS3A02G093400 | 59822956 | <u>1.44</u> | Xyloglucan endotransglycosylase/hydrolase protein 8-like | Cell wall-modifying enzyme that can loosen cell walls leading to cell expansion and elongation |
| TraesCS3A02G093500 | 59954598 | <u>65.07</u> | Xyloglucan endotransglycosylase/hydrolase protein 8-like | Cell wall-modifying enzyme that can loosen cell walls leading to cell expansion and elongation |
| TraesCS3A02G093600 | 60005281 | <u>0.51</u> | Xyloglucan endotransglycosylase/hydrolase protein 8-like | Cell wall-modifying enzyme that can loosen cell walls leading to cell expansion and elongation |
| **TraesCS3A02G093700** | 60028751 | <u>7.01</u> | **Xyloglucan endotransglycosylase/hydrolase protein 8** | Cell wall-modifying enzyme that can loosen cell walls leading to cell expansion and elongation |
| **TraesCS3A02G093800** | 60193220 | <u>0.47</u> | **Xyloglucan galactosyltransferase KATAMARI1 homolog** | Responsible for actin organization and the synthesis of cell wall materials |
| TraesCS3A02G094000 | 60203935 | <u>8.63</u> | Glycine-rich cell wall structural protein-like |  |
| **TraesCS3A02G094600** | 60446831 | *0.40* | **Cellulose synthase-like protein G2** | Cellulose biosynthesis and cell wall biogenesis |
| TraesCS3A02G096000 | 61311578 | *1.56* | Amino-acid permease BAT1 homolog | Involved in BR homeostasis. |
| TraesCS3A02G096200 | 61340198 | <u>1.00</u> | Protein NETWORKED 1A | Potentially couples different membranes to actin cytoskeleton |
| **QTL Marker** | **61343069** |  | **BS00003801** |  |
| **TraesCS3A02G096300** | 61346177 | <u>1.09</u> | **Protein NETWORKED 1A** | Potentially couples different membranes to actin cytoskeleton |
| TraesCS3A02G099400 | 64278873 | <u>2.87</u> | Serine/threonine-protein kinase BRI1-like 2 | Involved in brassinosteroid mediated signaling |

**Table 2.** List of candidate genes and mapping markers around the marker closest to the Hd QTL, BS00021976. Transcript name, start position (in CS sequence), expression level ratio between Avalon and Cadenza (ABS, <u>higher value</u> from Avalon, *higher value* from Cadenza), gene identity and gene function, if known.

| Gene ID | Start | ABS | ID | Function (if known) |
| --- | --- | --- | --- | --- |
| **Mapping marker** | 81477543 |  | **BS00022624** |  |
| TraesCS3A02G116300 | 84184377 | <u>1.01</u> | *TaGI* |  |
| **Mapping marker** |  |  | **wmc505** |  |
| TraesCS3A02G122600 | 97972874 | <u>1.84</u> | *Gibberellin 3-beta-dioxygenase 2-1* |  |
| **Mapping marker** | 106434896 |  | **BS00022148** |  |
| TraesCS3A02G133400 | 110332577 | <u>1.53</u> | *gibberellin 2-beta-dioxygenase 1-like* |  |
| TraesCS3A02G136500 | 114041077 | <u>1.05</u> | *SHAGGY-like kinase* |  |
| TraesCS3A02G143100 | 124172881 | <u>5.37</u> | *FT2* |  |
| TraesCS3A02G155200 | 147789276 | <u>1.03</u> | *IAA3* |  |
| TraesCS3A02G156500 | 151642540 | <u>1.11</u> | *ABI8* | ABA response |
| TraesCS3A02G159200 | 158467595 | *0.17* | *ARF1* | Auxin response |
| TraesCS3A02G164200 | 168837089 | <u>1.05</u> | Shaggy-related protein kinase | Brassinosteroid signalling pathway |
| **QTL Marker** | **168837302** |  | **BS00021976** |  |
| TraesCS3A02G167100 | 172629517 | <u>1.13</u> | Putative brassinosteroid receptor |  |
| TraesCS3A02G173300 | 191527521 | <u>1.06</u> | *Topless* |  |
| TraesCS3A02G176100 | 196761940 | <u>2.25</u> | flowering-promoting factor 1-like protein 1 |  |
| **Mapping marker** | 266691232 |  | **BS00110129** |  |
| **Mapping marker** | 310746858 |  | **barc19** |  |

### Gene candidates for Ht

A cluster of genes involved in cell-wall structure or synthesis, all of which could have a role in Ht form an interesting group around the QTL peak marker (Table 1). The clustering could suggest a close interaction between some or all of the genes. There are four, almost identical and apparently functional copies of a xyloglucan endotransglycosylase/hydrolase protein 8 gene in Avalon and Cadenza, except for one probably non-functional copy in Avalon. Networked 1A appears to have been duplicated at this locus as the two transcripts are quite different. The first copy is very well conserved between Cadenza and Avalon while the second copy shows a number of polymorphisms, which give both conservative and non-conservative amino acid changes in the Avalon protein compared to Cadenza and Chinese Spring. Other potential candidates are *cellulose synthase-like G2* (*CSL-G2*, involved in cellulose biosynthesis and cell wall biogenesis) and *BRI1-like 2 (BRI1L2*, involved in brassinosteroid mediated signaling). *CSL-G2* has four SNPs between Avalon and Cadenza within exons, resulting in two amino acid changes, while *BRI1L2* has no SNPs.

Transcriptomic analysis also shows expression level differences between some Avalon and Cadenza alleles in the cluster; most notably TraesCS3A02G093500, a Xyloglucan endotransglycosylase, showing increased expression in lines carrying the Cadenza allele with an ABS of over 65. KASP markers have been designed to SNPs within TraesCS3A01G093700.1 (Xg Copy 4), TraesCS3A01G093800.1 (Katamari), TraesCS3A02G094600 (CSL-G2), TraesCS3A01G096200.1 (Net1A_200). and TraesCS3A01G096300.1 (Net1A_300). Along with BS00022516 (co-segregating with BS00003801) these markers have been tested on the Watkins (Watkins Stabilised Collection of Hexaploid Landrace Wheats) and GEDIFLUX (Genetic Diversity Flux winter wheat collection) panels. The Gediflux panel gave three haplotypes in the region, only one of which shows recombination; this suggests the locus has already been fixed in this geographically and temporally more limited collection (Figure 6A). Analysis of Ht data from two trials of GEDIFLUX (2011 and 2016) indicate that both the Avalon and recombinant haplotypes are significantly different from the Cadenza haplotype with the recombinant being the shorter (Figure 6 A and B). The results suggest that the gene affecting Ht is at the distal end of the locus i.e *Xyloglucan endotransglycosylase* / Katamari / *cellulose synthase-like G2* or another unidentified gene.

**Fig 6.**
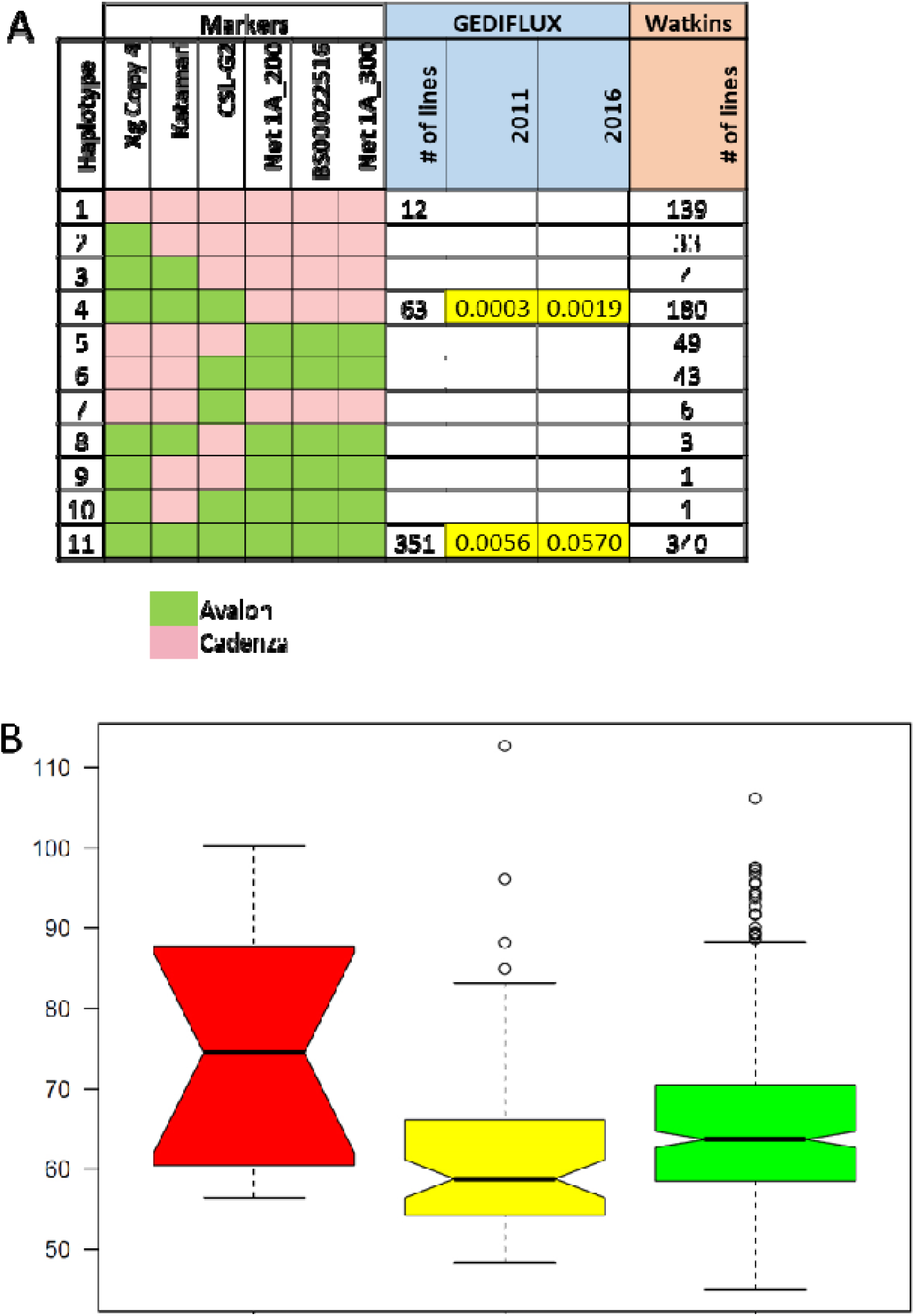
Haplotypes across the 3A Ht QTL locus in the GEDIFLUX and Watkins panels. A. Green and pink blocks represent Avalon or Cadenza at each marker for haplotypes 1-11. The significant differences between haplotype 1 and 4 or 11 are show for the GEDIFLUX 2011 and 2016 trials. B. Box plots indicating the Ht range for each haplotype (2011) data. Red is haplotype 1, yellow haplotype 4 and green haplotype 11 (from 6A.)

**Fig. 7.**
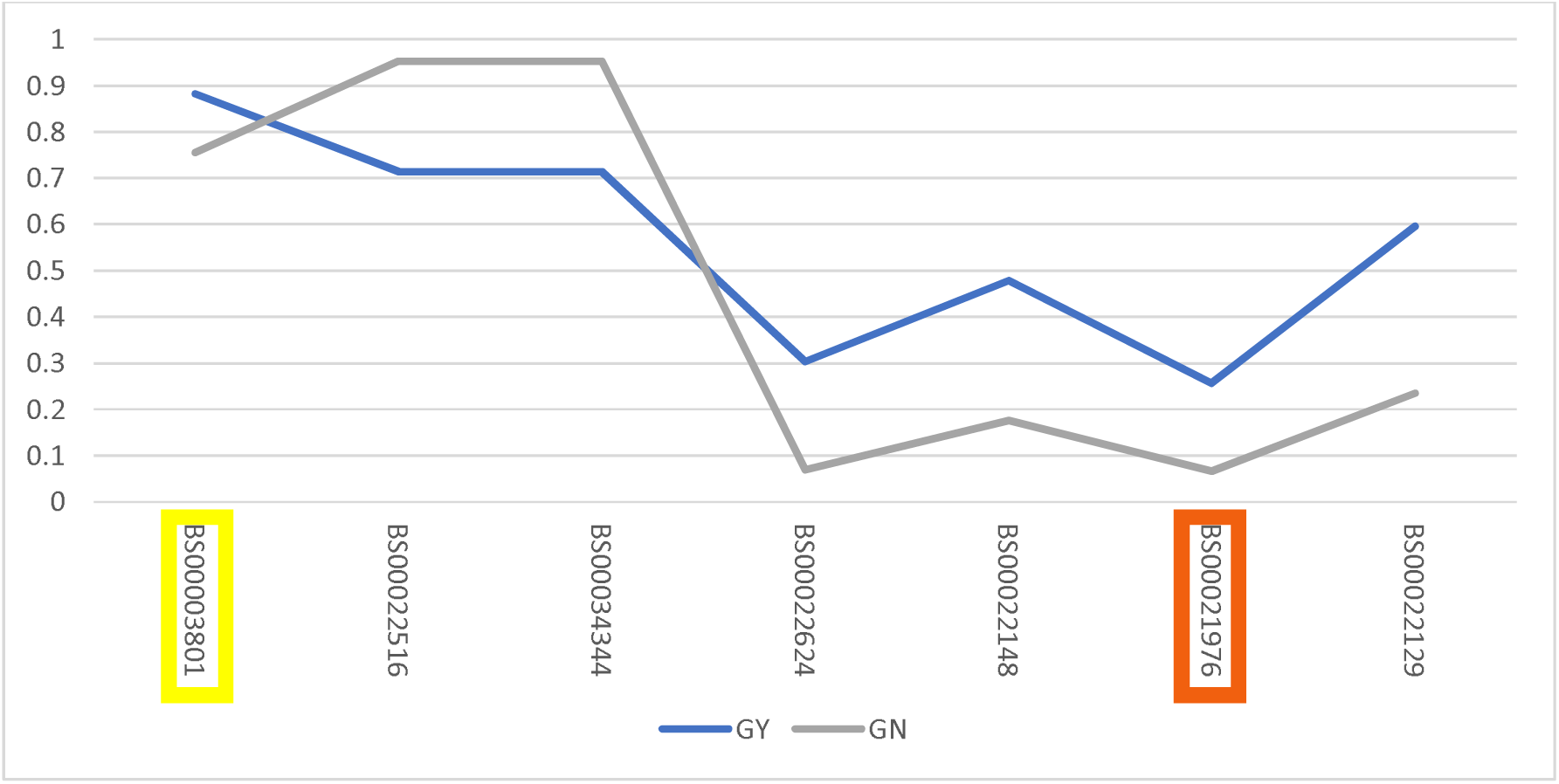
Single marker regression p values from H2016 yield component data plotted in the marker order of the wheat genome reference sequence. Peak marker for Ht (BS00003801) is boxed in yellow and for Hd (BS00021976) in orange.

The Watkins panel, a very diverse collection from across the globe, gave at least 11 haplotypes (Figure 6A) apparently due to multiple recombinantion events. Due to the difficulty in collecting accurate height data and the probability of population structure we cannot assign significance to the different haplotypes in Watkins. However the results suggest that this small region (less than 1.5 Mb) may be a hotspot for recombination allowing different combinations of genes which potentially affect plant height to come together. To examine this further we analysed the Axiom genotyping data of 4 Mb over the cluster from a larger panel of mainly hexaploid varieties (including the Watkins and GEDIFLUX lines). These data confirmed a high degree of recombination within the gene cluster (data not shown).

### Gene candidates for Hd

Prior to the publication of the IWGSC RefSeq v1.0 genome assembly from Chinese Spring (Appels et al., 2018) several genes involved with flowering and development were identified in the wider genetical vicinity of the Hd QTL. These genes are found in pathways known to be involved in flowering time: photoperiod (*GI* and *CDF1*), plant hormones i.e. gibberelin (*GA2ox-5-3* and *FPF1, Flowering Promoting Factor 1-like 1*), auxin (*ARF1* and *IAA3*), brassinosteroid signalling (*TaSK5*, a GSK3/SHAGGY-like kinase, and *BRI-associated receptor kinase*) and abscisic acid (*ABI8*). Publication of the IWGSC RefSeq v1.0 genome assembly allowed a closer examination of the genes around the marker at the Hd QTL peak. The marker with the highest association to the Hd QTL is KASP marker BS00021976. Table 2 shows possible gene candidates (based on gene function or expression differences) in the 60 Mb region either side of BS00021976, from its position in the CS sequence. We consider that *TaSK5, FT2* (*Flowering Locus T2*) or *FPF1* are the most promising gene candidates for Hd, based on likely gene function. BS00021976 maps at 169 Mb in the IWGSC RefSeq v1.0 genome assembly to a SNP from Cadenza in the 3’ UTR of TraesCS3A02G164200 (*TaSK5*). A KASP marker was developed to what appeared to be a second SNP in exon 4 of the same gene (in Avalon rather than Cadenza), causing an amino acid change. When this second marker was used for QTL mapping it did not associate with Hd and was mapped genetically more distally on 3AS. Examination of the Chinese Spring sequence at this position (102 Mb) indicates the presence of an unannotated GSK3/SHAGGY-like kinase pseudogene but we suggest this sequence must either incomplete or more likely that a duplicated copy of *TaSK5* is found at this location in Avalon as the CS sequences are not homologous enough to amplify the marker. In addition there is another SHAGGY-like kinase at 114 Mb, which appears to be functional but again has insufficient homology be amplified by the SNP2 marker. We therefore propose that the copy at 102 Mb has the exon 4 SNP, but in not involved in Hd, and the copy at 169 Mb, which has in 3’UTR SNP, can be considered a candidate, although it shows little difference in expression level between Avalon and Cadenza. In addition we suggest there has been a rearrangement around the Hd QTL as there is a divergence in the expected order of markers from the CS sequence when arranged according to genetic rather than physical distance (see Figure 5). *FT2* is approximately 4.3 kb, consisting of four small exons and a large intron 2 of 2.9 kb, the sequence of which is incomplete. With the available data three SNPs have been identified between Avalon and Cadenza, in the few transcripts from intron 2, but there are no polymorphisms in the coding regions. RNAseq data and visualization in IGV indicates that most of exon 1 and all of exons 2 and 3 are missing in the Cadenza transcript at this developmental stage. The missing exons contain the majority of the PEBP domain and would therefore affect FT2 function in Cadenza. Expression of the gene is higher in Avalon and higher expression of *FT2* has been shown to cause earlier flowering in (Shaw *et al.*, 2019).

*FPF1* does not show any polymorphisms between Avalon and Cadenza but is more highly expressed in Avalon than Cadenza and in Arabidopsis is expressed in the equivalent of the sowing -> DR -> TS growth stages in wheat (ref).

### Effect of 3A Ht and Hd loci on grain yield

Eight informative recombinants were grown together with 3A NIL controls and the parents Avalon and Cadenza over two field seasons. As these lines were grown in relatively large (6 m^2^ plots) this provided the opportunity to dissect the locus in terms of grain GY. In harvest year 2016 and 2017 experiments the Ht and Hd results reported here and in Farre et al (2016) were repeated with Cadenza alleles at the 3A locus resulting in a crop which was later heading (by one day) and taller (11.7cm). The following season the direction of allelic effect stayed the same for these two traits but was reduced to 0.67 days for Hd and 10.2 cm for Ht. For GY, the Cadenza allele conferred an increase in H2016 which was driven by grains per unit area, as was the case for Farre et al (2016), but there were no GY difference between the NILs for H2017, for this reason the effect of the locus on GY was only analysed using H2016 data. In Figure 8 p values of each of the markers spanning the locus are shown for GY, TGW, and GN. None of these differences show >0.05 p values. However, a trend of increasing additive effect associated with Cadenza alleles (not shown) and decreasing p value is seen for markers closer to the Hd locus. Single marker regression showed that *XBS00021976* was most significantly associated with grains per unit area (p value 0.17) and GY (p value 0.28).

## Discussion

In the grasses stem extension and reproductive development (and so the formation of yield components) are very tightly linked developmental processes. Stem extension begins once the inflorescence has reached terminal spikelet and ends around the same time as anthesis, so there is correlation between traits associated with each process. This correlation can be seen at the molecular level with the same gene affecting stem extension, phenology, and GY, for example *Ghd8* in rice (Yan *et al.*, 2011). However, we show that the Ht and Hd QTL collocated on chromosome 3A (Griffiths *et al.*, 2009; Griffiths *et al.*, 2012; Ma *et al.*, 2015) are genetically linked but independent effects. We have used genetic recombination to separate them, effectively developing sets of sub-NILs which display height and heading differences in isolation. This opens the way for the independent selection of these loci in breeding. Analysis of the haplotypes across the Ht locus within the AE Watkins landrace collection and the Gediflux collection of 20^th^ Century European winter wheat, show that historical recombinants of this kind have occurred multiple times with 11 haplotypes apparent. The genetic separation of Ht and Hd also allowed us to ask how each of them was contributing to the GY QTL which we had previously shown to collocate on chromosome 3A. Our data shows that GY is a pleiotropic effect of the Ht locus but the proposition that the GY and Hd increasing alleles of Cadenza belong to the same gene is plausible.

### Gene candidates for Ht, Hd, and GY

The Hd interval spans the centromere of chromosome 3A over a relatively large physical distance. The equivalent region of Chinese Spring is 229 Mb and contains 536 genes. This part of the 3A locus was intractable to further genetic resolution in this work due to reduced levels of recombination encountered in centromeric regions. The 3A Ht effect is more distally located on 3AS in a 5 Mb equivalent region of Chinese Spring containing 146 genes. In spite of these large gene numbers analysis of known function, polymorphism between Avalon and Cadenza, and expression level differences did support the proposal of gene candidates for the 3A QTL complex that we had set out to dissect.

For Ht the cluster of cell wall related genes and expression level differences from them could be resolved to some extent using the historical recombination events present in the Gediflux collection. This pointed towards the *Xyloglucan endotransglycosylase*, Katamari, and *cellulose synthase-like G2* part of the cluster being prioritised as most likely containing the causative gene/s. Xyloglucan is an essential component in the formation and function of the plant cell walls and both the xyloglucan endotransglycosylases and Katamari (MUR3) are involved in these processes. Xyloglucan endotransglucosylases/hydrolases catalyze the endo cleavage of xyloglucan polymers and appear to have a role in cell wall restructuring. The rice orthologue, *OsXTH8*, is thought to be involved in this process and is regulated by GA, with increased expression leading to increased plant height (Jan *et al.*, 2004). Plants synthesise xyloglucan which contain galactose in two types of side chain. In Arabidopsis mutants of MUR3 missing one type of side chain have a dwarf phenotype (Kong *et al.*, 2015). Cellulose is the most important component of plant cell walls, required to maintain shape and rigidity. Cellulose-synthase like genes may be involved in the synthesis of the back-bones of hemicelluloses In rice DNL1 is a major QTL for plant height and encodes OsCSLD4 (Ding et al 2015). An unidentified glycine-rich cell wall structural protein-like is also located within this region.

For Hd and GY it is *FT2* that stands out as a candidate in terms of it’s known dual effects on Hd and spike fertility. As reported by (Shaw *et al.*, 2019) in durum wheat *FT2* late alleles also confer increased spikelet number which was also observed by (Farré *et al.*, 2016), in the same material described here. In addition (Shaw *et al.*, 2019) observed floret number per spikelet increases also observed by (Ochagavía *et al.*, 2018) in the Avalon/Cadenza NILs. In (Shaw *et al.*, 2019) the observed fertility changes were not accompanied by a yield increase but they do provide a physiological footprint, beyond simple phenology, that are analogous to spike fertility, and GY effects reported here and in our previous work.

### Further delimitation of the GY QTL

Most of the Ht and Hd work described here was done in small plots which are not appropriate for yield estimation. However, informative recombinants were grown in large (6 m^2^) replicated plots to better understand how the independent Ht and Hd effects influence GY. This experiment did not identify significant GY differences but did show a clear trend for a yield difference, equivalent to that seen between NILs, in recombinants differing for the Hd QTL but the same mean GY for Cadenza and Avalon alleles at the Ht QTL. The evidence presented here supports the location of the GY effect as within the newly defined Hd locus. It is interesting to note that in the two seasons of yield trial described here, the year when the NILs parents did not significantly differ in Hd they did not differ in GY, while the Ht phenotype was strongly expressed. As already stated, FT2 is a candidate for the Hd and GY QTL. The work presented here supports the value of further exploration of the role of FT2 on spike fertility and GY.

### Environmental sensitivity of Hd effect

It is important to understand how the 3A Hd effect interacts with environmental signals, not only for it’s role in adaptation but also our speculation that the same gene is influencing GY. Controlled environment studies showed that this is not a new *Vrn* or *Ppd* gene. In fact it fulfils the classical categorisation as earliness *per se* (*eps*), with a difference in Hd still evident after saturating vernalization and photoperiod. It is also clear that the *eps* effect acts on the vegetative to floral transition. This QTL does exhibit interesting genotype by environment interactions. We showed that, when expressed, the developmental difference is apparent at the vegetative and early reproductive phases, with no difference in the duration of the late reproductive phase. The Hd effect was not expressed after spring drilling. In the vernalization experiments the additive effect increased when vernalization was fully satisfied. Elsewhere (Ochagavia et al 2018) showed that the NILs did not show any phenology differences in Northern Spain. In experiments not presented here we also showed no significant effect on phenology at 16 and 21°C controlled environment experiments. Taken together, these data point towards the likely role of ambient temperature sensitivity differences, during vegetative growth, between Avalon and Cadenza alleles. Previous studies challenged the idea of earliness *per se* as a wholly autonomous developmental process occurring without environmental interaction seemed unlikely. Indeed, we showed that *EPS-D1* exhibits a strong interaction with temperature (Ochagavía *et al.*, 2019) and it seems likely that this is also the case the 3A Hd effect described here, that should probably be named *EPS-A1*.

## Supporting information

Meteorological data

3A locus transcript analysis

## Supplementary data

Fig. S1. Meteorological data relating to field experimentation.

Table S1. Expression level differences for predicted genes of whole 3A locus

## Acknowledgements

Funding from Defra, BBSRC and AHDB towards the LINK project “Exploiting novel genes to improve resource use efficiency in wheat (BBS/E/J/000CA399)” is gratefully acknowledged as we as the Defra Wheat Genetic Improvement Network and the BBSRC funded Designing Future Wheat Institute Strategic Programme.

## Abbreviations

BSA: Bulk Segregant Analysis
CIM: Composite interval mapping
DR: Double ridge
*eps*: Earliness *per se*
NIL: Near Isogenic Line
Ppd: Photoperiod
QTL: Quantitative Trait Locus
Rht: Reduced height
RPKM: Reads Per Kilobase of transcript per Million mapped reads
S: Sowing
TS: Terminal spikelet
Vrn: Vernalization

