## Supplementary material for "Resolving a QTL complex for height, heading, and grain yield on chromosome 3A in bread wheat": Meteorological data

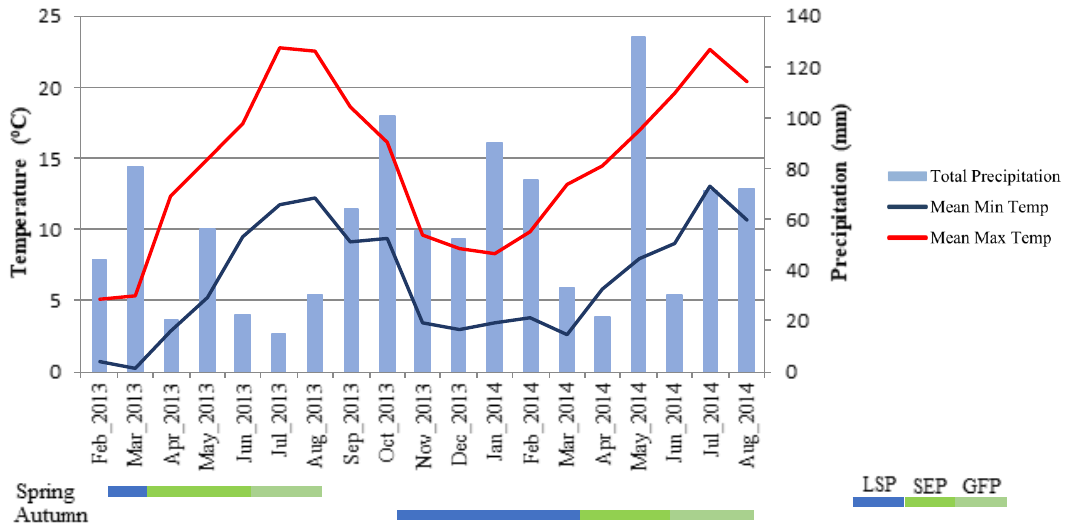

Figure 1.      Means of minimum (blue line) and maximum (red line) temperatures and cumulated rainfall (blue bars) during the months of the growth cycle of the spring and autumn sown trials (LSP = period from seedling emergence to first node detectable, SEP = period from first node detectable to anthesis and GFP = period from anthesis to physiological maturity).
